# CanLncG4: A database curated for the assessment of G4s in the lncRNAs dysregulated in various human cancers

**DOI:** 10.1101/2024.02.21.581359

**Authors:** Shubham Sharma, Muhammad Yusuf, Noman Hasif Barbhuiya, Harshit Ramolia, Chinmayee Shukla, Deepshikha Singh, Bhaskar Datta

## Abstract

Long non-coding RNAs (lncRNAs) comprise a substantive part of the human genome and have emerged as crucial participants of cellular processes and disease pathogenesis. Dysregulated expression of lncRNAs in cancer contributes to various hallmarks of the disease, presenting novel opportunities for diagnosis and therapy. G-quadruplexes (G4s) within lncRNAs have gained attention, though their systematic evaluation in cancer biology is yet to be performed. In this work, we have formulated CanLncG4, a comprehensive database integrating experimentally validated associations between lncRNAs and cancer, and detailed predictions of their G4-forming potential. CanLncG4 categorizes predicted G4 motifs into anticipated G4 types and offers insights into the subcellular localization of the corresponding lncRNAs. It provides information on lncRNA-RNA and lncRNA-protein interactions, together with the RNA G4-binding capabilities of these proteins. To ensure the accuracy and validity of the data sourced from various databases, a meticulous examination of the output data was conducted to identify any discrepancies, including incorrect, missing, or duplicate entries. Additionally, scientific literature mining was performed to cross-validate the gathered information. Data from G4-prediction tools was generated using multiple parameter combinations to determine the parameters that yield more relevant and accurate predictions of the G4-forming potential. We validate our *in silico* G4-prediction pipeline through *in vitro* experiments, affirming the presence of G4s within specific cancer-dysregulated lncRNAs, thereby illustrating the predictive capability of CanLncG4. CanLncG4 represents a valuable resource for investigating G4-mediated lncRNA functions in diverse human cancers. It is expected to provide distinctive leads about G4-mediated lncRNA-protein interactions. CanLncG4 comprehensively documents 17,666 entries, establishing correlations between 6,408 human lncRNAs encompassing their transcript variants, and 15 distinct types of human cancers. The database is freely available at https://canlncg4.com/, offering researchers a valuable tool for exploring lncRNA and G4 biology towards cancer diagnosis and therapeutics.

## Introduction

Only a fraction of the human genome directly codes for proteins, while the majority is composed of non-coding regions. Considering the complexity of the human genome, this apparent paradox has been partially understood by the discovery of non-coding RNAs (ncRNAs) that play crucial regulatory roles in cells.^1–5^ Among ncRNAs, long non-coding RNAs (lncRNAs) with lengths of 200 or more nucleotides are a prominent subset, orchestrating diverse functions in cellular physiology and disease progression.^6, 7^ LncRNAs are characterized by tissue-specific and spatiotemporal expression patterns, and deploy complex three-dimensional structures towards cell growth, differentiation, and development.^8^ The dysregulation of lncRNAs in various diseases, including cancer, diabetes, autoimmune disorders, and neurodegenerative conditions, highlights their deep connection with cellular homeostasis and the potential for innovative therapeutic interventions.^9^

Cancer remains a prominent cause of disease-related mortalities worldwide due to ineffective diagnostic, prognostic, and therapeutic regimens. The molecular complexities underlying carcinogenesis and cancer progression, together with the emergence of resistance mechanisms and long-term side-effects of treatment strategies, pose challenges for the effective cure of the disease.^10, 11^ LncRNAs are dysregulated and mutated across various cancer types and can exhibit oncogenic or tumor-suppressing functions.^11–16^ The dysregulation of lncRNA disrupts cellular homeostasis and contributes to several aspects of carcinogenesis, such as cancer cell proliferation, migration, invasion, metastasis, apoptosis, altered cell metabolism, cell cycle regulation, and treatment resistance.^17–22^ The tissue-specific expression of lncRNAs, coupled with their detectability in bodily fluids, positions them as promising candidates for cancer biomarkers.^23, 24^ Our understanding of lncRNAs in the context of cancer remains nascent compared to other RNA counterparts such as miRNAs. Key questions persist regarding the precise mechanism of lncRNA activity vis-à-vis cancer pathogenesis.^16, 25^

RNA molecules are capable of folding into a plethora of secondary and tertiary structures, ranging from duplexes, stem-loops, and simple hairpins to complex motifs like bulges, triplexes, and G-quadruplexes (G4s). These structural variations are correlated with RNA functionality, with dynamic changes often profoundly impacting the functional outputs.^5^ G4s are formed by G-rich sequences with characteristic sequence traits, and are stabilized by monovalent cations under physiological conditions.^26^ RNA G4s are prevalent in human mRNAs, telomeric RNA, and viral RNA genomes, and are purported to participate in transcription termination, pre-mRNA processing, mRNA targeting, translation regulation, and telomere maintenance.^27, 28^ Within the cellular milieu, RNA G4s exist in a dynamic equilibrium with their unfolded counterparts. This equilibrium can be influenced by factors like transcription rates and the presence of RNA-binding proteins possessing chaperone and helicase activities.^29^ G4s have emerged as key players in the molecular mechanisms underlying cancers, neurological disorders, and infectious diseases, highlighting their significance in disease progression and, hence, their potential as therapeutic targets.^30–37^

G4s are a prominent higher-order structure within the repertoire of structures adopted by lncRNAs.^5^ Analysis conducted across the transcriptome using a G4-prediction tool, Quadfinder, has revealed notable putative G4 motifs within lncRNAs typically ranging in size from 200 to 300 bases and characterized by short loops.^38^ Although the precise biological roles of G4s within lncRNAs remain relatively unexplored, a handful of reports have shed light on their significance in select lncRNAs, including *hTERC*, *GSEC*, *RPPH1*, *XIST*, *RMRP*, *REG1CP*, *NEAT1*, *LUCAT1*, *TERRA*, and *NORAD*.^25, 29, 39–45^ Recent investigations have delved explicitly into the role of G4s within the *MALAT1* lncRNA, elucidating their involvement in the regulation of cellular functions.^46, 47^ These studies also underscore the participation of G4s and a spectrum of G4-binding proteins (G4BPs) in mediating the functions of lncRNAs, further emphasizing the interplay between G4s and regulatory proteins.

While dysregulation of lncRNAs has been implicated in diseases like cancer, there remains a need for a comprehensive investigation into the precise impact of lncRNA secondary structures and their influence on protein interactions to regulate disease progression.^45, 48^ Among the subset of lncRNAs harboring G4s, *NEAT1*, *H19*, *CRNDE*, and *HOTAIR* have emerged as noteworthy contributors to cancer development, underscoring the pivotal role of these structures in fostering cellular malignancy.^41, 48–50^ Recent studies focusing on *GSEC*, *REG1CP*, and *LUCAT1* lncRNAs, which exhibit upregulation in colorectal cancer, further highlight the significance of lncRNA G4-protein-mediated regulation of gene expression in the pathogenesis of cancer.^25, 40, 42, 45^

The early *in silico* methods for predicting putative G4s in DNA and RNA relied solely on biophysical experiments. However, with advancements in genome sequencing technologies, sequencing data has become reasonably simple, easily accessible, and cost-effective, enabling the development of sequence-based computational prediction tools.^51^ Some of the G4-motif prediction tools that can predict regions with high G4-forming propensity in DNA and RNA, such as Quadfinder, Quadparser, Quadruplexes, AllQuads, and ImGQfinder, are based on regular expression matching.^51–55^ Other tools, such as QGRS Mapper, pqsfinder, G4P calculator, cG/cC, and G4Hunter, are based on the score-based ranking of the putative sequences to enable prediction of the most probable G4-forming sequence.^56–60^ Some other tools, such as the G4RNA screener and Quadron, are based on machine learning methods.^61, 62^ A select number of resources focus specifically on RNA G4s, including the databases G4LDB and G4IPDB which catalog RNA G4-ligands and proteins, and resources like ONQUADRO and DSSR-G4DB which display RNA and G4 structures.^63–66^ Additionally, databases like G4RNA, GRSDB2, GRS_UTRdb, and QUADRatlas provide information on experimentally validated and predicted RNA G4s, particularly in mRNAs and untranslated regions (UTRs).^67–69^ A limited number of resources have systematically documented lncRNAs in cancers. The platform Lnc2Cancer focuses on lncRNA-cancer associations, while databases like NPInter and LncTarD catalog lncRNA interacting partners and disease associations.^70–72^ Despite the availability of these informatics and computational tools, there is an absence of platforms that integrate different data types and enable the correlation of G4s with established lncRNAs, and their associations with cancer. Such platforms could provide users with detailed information on lncRNA-cancer associations, lncRNA G4-prediction, lncRNA subcellular localization, and lncRNA-G4 interacting partners. Additionally, these platforms could facilitate the exploration of G4-mediated lncRNA-protein interactions, thereby enabling the discovery of cellular mechanisms involving lncRNA G4s.

In this work, we introduce CanLncG4, a meticulously curated database consolidating experimentally validated associations between lncRNAs and a large number of human cancers. CanLncG4 offers an extensive G4-prediction analysis for each lncRNA transcript variant, leveraging well-established tools like QGRS mapper and G4Hunter. Additionally, it categorizes the PQS predicted by these tools into anticipated G4 types (2G, 3G, and 4G), providing valuable structural insights into the potential G4 motifs. CanLncG4 can compare PQS predicted by the established G4-prediction tools and facilitates the re-analysis of catalogued lncRNA sequences by modifying analysis parameters. The database incorporates G4-prediction tools as standalone features, allowing users to comprehensively assess, classify, and compare G4-predictions linked to any specified sequence and NCBI accession ID. CanLncG4 offers insights into the subcellular localization of catalogued lncRNAs across a diverse range of cell lines. The database conducts a comprehensive meta-analysis of lncRNA interacting partners, utilizing data from NPInter v4.0 and LncTarD. Moreover, it provides information about the established RNA G4-binding capabilities of proteins interacting with the catalogued lncRNAs, drawing from QUADRatlas, G4IPDB, and scientific literature mining. Considering that the majority of catalogued lncRNAs contain putative G4-forming regions, details of proteins interacting with these lncRNAs, and their established RNA G4-binding potential, can serve as an informed starting point for investigating G4-mediated lncRNA-protein interactions. We corroborated the *in silico* G4-assessment pipeline of CanLncG4 through *in vitro* experiments on high-scoring PQS derived from randomly selected cancer-dysregulated lncRNAs. Our results indicate the presence of G4s within these lncRNAs, thereby affirming the predictive capability of our methodology. CanLncG4 serves as a comprehensive platform for accessing information about cancer-dysregulated lncRNAs, their G4-forming potential and interacting partners. CanLncG4 is freely accessible at https://canlncg4.com/.

## Material and Methods

### Data collection and generation

#### A. *In silico* G4-prediction in cancer-dysregulated lncRNAs

The comprehensive list of lncRNAs dysregulated in diverse human cancers, their expression patterns, methodology of identification, and PubMed ID of research articles used to obtain these details, were obtained from Lnc2Cancer 3.0 (Figure 1A).^70^ The aliases of all these lncRNAs were manually compiled from GeneCards.^73^ The NCBI accession numbers and the corresponding FASTA sequences of all the identified lncRNAs and their functional transcript variants with validated and reviewed RefSeq status were retrieved from NCBI Nucleotide, a database comprising DNA and RNA sequences from numerous resources such as RefSeq, GenBank, TPA, and PDB.^74^ A meticulous examination of the data sourced from the mentioned databases was conducted to identify discrepancies such as incorrect, missing, or duplicate entries, and to ensure the accuracy and validity of the data. Additionally, the gathered information was cross-validated with available scientific literature. CanLncG4 documents 17,666 entries establishing correlations between 6,408 human lncRNAs including their transcript variants, and 15 distinct types of human cancers. The cancers that are included in CanLncG4 version 1.0 are General: Head and neck, skin, lung, liver, gastric, colorectal, brain, bone, blood; Male-dominant: prostate and testicular; Female-dominant: breast, ovarian, uterine, and cervical cancer.

**Figure 1.**
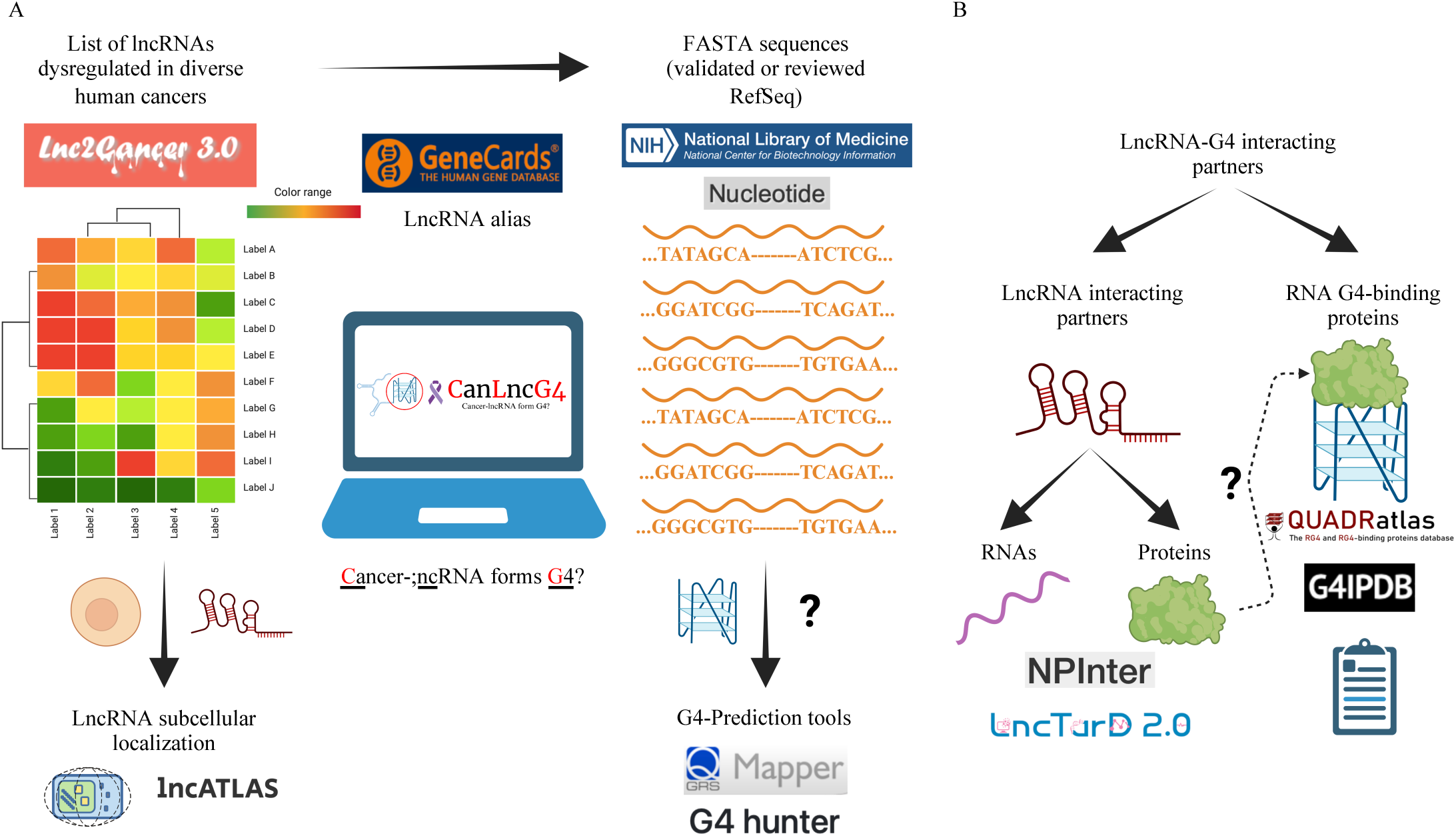
Schematic of CanLncG4 workflow. A) *In silico* prediction of G4 formation in the lncRNAs dysregulated in human cancers using Lnc2Cancer 3.0: lncRNA-cancer association, GeneCards: lncRNA aliases, NCBI Nucleotide: lncRNA FASTA sequence; G4-prediction tool: QGRS mapper and G4Hunter; LncATLAS: lncRNA subcellular localization. B) Identification of lncRNA-G4 interacting partners using NPInter v4.0 and LncTarD 2.0: LncRNA-RNA and LncRNA-Protein interactions; QUARRatlas, G4IPDB, and scientific literature mining: RNA G4 binding proteins (RGBP).

For identification of PQS within these lncRNAs, their FASTA sequences are imported to the QGRS mapper, a tool that presents the data on the constitution and distribution of Quadruplex-forming G-rich sequences (QGRS) (Figure 1A). The QGRS mapper identifies the sequences as PQS based on the canonical sequence of the format: G_x_N_y1_G_x_N_y2_G_x_N_y3_G_x_, where x = number of G-quartets in the G4/ single G-tract (tandem repeats of guanines) length and y1, y2, y3 = loop lengths, which collectively define the G-Score of each PQS.^56^ All the PQS possible in catalogued lncRNAs and their transcript variants are identified using the following parameters: max length: 45; min G-group: 2; loop size: 0 to 36. PQS with higher G-Scores have a greater propensity to form G4s. However, to ease the PQS selection and streamline the user experience, only the highest-scoring PQS amongst all the overlapping ones are presented.

The PQS within these lncRNAs are also identified using G4Hunter, a tool for identifying putative G4-forming motifs in nucleic acids based on the G-richness and G-skewness of the query sequence (Figure 1A).^60^ The FASTA sequences of each catalogued lncRNA and their transcript variants are fed into the G4Hunter algorithm, and all the possible PQS are identified using the following parameters: window size: 45; threshold: 0.9. The output G4H Score conveys the probability of G4 formation by the query sequence. While exploring the G4Hunter tool, complexity in terms of the PQS presented by the G4Hunter can be encountered. In addition to presenting all the overlapping PQS and their respective scores as per the set parameters, a consensus sequence containing all the overlapping PQS but with a different score is also presented. This PQS representation strategy is followed for all the PQS identified by the G4Hunter, which results in a massive volume of output data that may overwhelm or distract the user, eventually preventing them from selecting the most promising candidate. Accordingly, the G4Hunter algorithm is slightly modified to present only the highest scoring PQS amongst the overlapping ones. Multiple parameter combinations were employed to identify PQS within these lncRNAs using QGRS mapper and G4Hunter. The previously mentioned parameters yielded more relevant and accurate predictions of the G4-forming potential.

The cytoplasmic to nuclear localization: relative concentration index, expression values, and distribution plots for the catalogued lncRNAs across diverse cell lines were sourced from the LncATLAS database to describe their subcellular localization (Figure 1A).^75^

#### B. Identification of LncRNA-G4 Interacting Partners

The details of experimentally validated RNA and protein interacting partners of catalogued lncRNAs were sourced from NPInter v4.0 and LncTarD 2.0.^71, 72^ The information regarding the experimentally validated RNA G4-binding proteins (RGBP) interacting with the catalogued lncRNAs were obtained from QUADRatlas, G4IPDB, and scientific literature mining (Figure 1B).^64, 69, 76^

### Architecture of Web Application

A detailed overview of the technology stack and architectural decisions underlying the web application hosted at https://canlncg4.com/ is provided in this section. It highlights the key components and methodologies employed to optimize both the user experience on the frontend and the efficiency of the backend operations.

#### A. Frontend Architecture

The web application hosted at https://canlncg4.com/ employs a technology stack designed to optimize both the user experience and backend efficiency. The frontend architecture is built on Next.js (v13.4), a React framework known for its server-side rendering capabilities. This choice ensures that pages are pre-rendered on the server, leading to faster load times and improved SEO. The frontend is deployed on Vercel, a platform that aids in hosting for JAMstack architectures, providing automatic scaling and a global CDN for enhanced accessibility and speed. In this connection, responsiveness and user experience are central to the frontend design, offering a comprehensive suite of tools and controls that facilitate efficient database querying via the GUI.

#### B. Core Backend Architecture

The application’s backend infrastructure exhibits a multi-tiered approach. The primary backend is developed using Python Flask (v3.0.1), a micro web server framework. This backend is responsible for the core logic and data processing of the application and is hosted on an Ubuntu (LTS 22.04.02) server on an AWS (Amazon Web Services) EC2 (t2.micro) instance. A distinctive feature of the backend architecture is its approach to data management. Instead of relying on traditional SQL or NoSQL databases, the application utilizes the pandas (v1.4.0) library to read data files stored in Excel format and create an in-memory SQLite database. This approach ensures data is readily available in RAM, leading to significantly faster query execution times. As the data resides in memory, there is no need for additional caching mechanisms to achieve high throughput, simplifying the overall architecture. Moreover, as we do not expect any writes to our databases, we utilize the same database connection across multiple threads, leading to a lower memory footprint.

The Ubuntu server hosting the Flask application is configured with Nginx (v1.18) acting as a reverse proxy. This setup enhances security by shielding the Flask server from direct external access and improves load handling and distribution, which may be a case for the future. We configured the security group and firewall for the backend server on AWS to only allow requests from the Vercel. Notably, despite Vercel’s support for serverless Python functions, we consciously avoided deploying the Flask server on Vercel. This decision was driven by concerns over cold start delays and the potential for exploitation through the unintended or malicious invocation of serverless functions.

#### C. Connectivity of Frontend and Backend

Communication between the frontend and backend is streamlined through a REST API server, which is implemented in Node.js and deployed on Vercel. This server serves as an intermediary layer, effectively enabling efficient data exchange between the Next.js frontend and the Flask backend. By functioning as a bridge, this intermediary layer fully decouples the frontend, the API server, and the core logic components.

### Oligonucleotides

We randomly selected 5 cancer dysregulated-RNA sequences having only one functional transcript variant from CanLncG4 (Table S1). The RNA sequences were prepared by *in vitro* transcription (IVT) of cDNA templates. The cDNA antisense template and T7 RNA promoter sense strand were designed with SnapGene software (Dotmatics, www.snapgene.com, USA). The DNA oligonucleotides were procured from Sigma-Aldrich Chemicals Pvt. Ltd., Bangalore, India. All oligonucleotides were reconstituted in nuclease-free water to a final concentration of 100 µM and stored at −20 °C until further use. Details regarding the sequences of the DNA oligonucleotide mentioned above are accessible in Table S1.

### Reagents

The DNase I (RNase-free) (Catalog no. M0303L), Monarch^®^ RNA Cleanup Kit (Catalog no. T2050L), HiScribe^™^ T7 High Yield RNA Synthesis Kit (Catalog no. E2040S), and RNase Inhibitor, Murine (Catalog no. M0314L) were procured from New England Biolabs Pte. Ltd., Singapore. TMPyP4 (Catalog no. 613560) and Thioflavin T (Catalog no. T3516) were obtained from Sigma-Aldrich Chemicals Pvt. Ltd., Bangalore, India. All reagents were prepared following the manufacturer’s protocol and stored under the recommended conditions.

### Preparation of RNAs

The templates for IVT were designed using an approach that we have recently communicated. Briefly, the IVT templates comprised the following components: (1) antisense sequences of transcription initiator, (2) cDNA of PQS corresponding to the RNAs, (3) an extra 30nt long sequence, towards the 5’-end of the antisense sequence of the T7 RNA promoter. The conventional transcription initiation sequence CCCTTT was replaced with CGCTTT in the cDNA antisense templates to lower heterogeneity and preclude the mis-incorporation of additional G-tracts to the existing PQS. The resuspended sense strand oligonucleotide (T7 RNA promoter) was mixed in an annealing buffer in equimolar proportion to the cDNA antisense template oligonucleotide (Table S1). The annealing buffer was 10 mM Tris-Cl (pH 7.5) with 50 mM NaCl and 1 mM EDTA (pH 8.0). Annealing of samples was performed by heating at 95 °C followed by spontaneous cooling to room temperature. The absorbances of annealed oligonucleotides at 260 nm were obtained using a NanoDrop™ 2000c spectrophotometer (Thermo Fischer Scientific India Pvt. Ltd., Mumbai, India), to determine their concentration. These absorbances at 230 nm, 260 nm, and 280 nm were used for the estimation of A_260/280_ and A_260/230_ values as measures of purity of the annealed oligonucleotides. IVT using the annealed oligonucleotides was performed using the HiScribe™ T7 High Yield RNA Synthesis Kit, following the manufacturer’s protocol. Degradation of the DNA template oligonucleotides was effected by treatment with DNase I (RNase-free) followed by purification by Monarch® RNA Cleanup Kit. The purified RNAs were reconstituted in nuclease-free water and supplemented with RNase inhibitor (Murine). The concentration and purity of purified RNAs were estimated using a NanoDrop™ 2000c spectrophotometer. Folding of the IVT RNAs into G4s was performed by heating at 95 °C for 5 minutes in the presence of a folding buffer: 10 mM Tris-Cl (pH 7.5) and 0.1 mM EDTA (pH 8.0), supplemented with or without the monovalent cations (KCl or LiCl), followed by spontaneous cooling to room temperature.

### Circular Dichroism Spectroscopy

Circular dichroism (CD) spectra for the RNAs (5 µM) folded in the presence or absence of 100 mM KCl/ LiCl were recorded on a J-815 CD Spectropolarimeter (JASCO Inc., USA). Spectra were recorded between 220 – 320 nm 25 °C. The parameters used for CD spectral measurements were as follows: data pitch: 1 nm; sensitivity: standard; DIT: 1 sec; bandwidth: 0.50 nm; scanning speed (100 nm/ min), and accumulations: 3. CD measurements on samples with TMPyP4 (meso-5,10,15,20-Tetrakis-(N-methyl-4-pyridyl) porphine) were performed by titrating folded RNAs (5 µM) with TMPyP4 (5 µM) followed by heating the mixture at 37 °C for 10 minutes. CD spectra were recorded after each titration until a final concentration of 25 µM was reached. Spectral data were smoothed with the Savitzky-Golay method and 20 points of window. Mean CD (dmeg) values for respective RNAs were plotted against the wavelengths using the Origin (Pro), Version 2017 (OriginLab Corporation, USA).

### Thioflavin T (ThT) Fluorescence enhancement assay

ThT fluorescence enhancement assay was performed on RNAs (2 µM) folded in the presence or absence of 100 mM KCl/ LiCl. Samples were mixed with ThT (2 µM) and the ThT fluorescence spectra were recorded using a BioTek Cytation Hybrid Multimode Reader (Agilent Technologies International Pvt. Ltd., Manesar, India). The excitation (400 – 460 nm) and emission (470 – 650 nm) spectra, along with endpoint fluorescence emission, were recorded with emission and excitation at 488 nm and 445 nm, respectively. The data were recorded in triplicates across two independent studies. The mean fluorescence intensities (arbitrary units) of excitation and emission spectra were plotted against the wavelength after smoothening the data with the Savitzky-Golay method and 20 points of window using Origin (Pro). The ThT fluorescence fold-enhancement was calculated from the endpoint ThT fluorescence in the presence of RNA/ absence of RNA: F/ F_0_. The mean ThT fluorescence fold-enhancement was plotted with standard error of mean against different monovalent cations used for the respective RNAs, using GraphPad Software (Dotmatics, USA). Ordinary one-way ANOVA was employed for the statistical analysis, and the resulting statistical significance (P-values) are denoted with asterisks (*) in GP style: 0.0332 (*), 0.0021 (**), 0.0002 (***), <0.0001 (****). Non-significant P-values are not represented.

## Results and Discussion

### Workflow of *in silico* G4-prediction in cancer-dysregulated lncRNAs

#### A. LncRNA-cancer associations

CanLncG4 is projected as a platform that presents experimentally validated associations between lncRNAs and various human cancers, while predicting their G4-forming potential (Figure 1A).^70^ The general search bar on the home page can be used to search the database by the lncRNA or the cancer name (Figure S1 A). The search output presents the lncRNA and cancer name, lncRNA expression pattern, methodology of identification, PubMed ID of research articles, details of G4 prediction, details of subcellular localization, and lncRNAs alias, with regards to the searched lncRNA or the lncRNAs associated with the searched cancer. While the PubMed ID can be selected to navigate to the respective research article, the details of G4 prediction and subcellular localization can be selected to navigate to the appropriate section (Figure S1 B, C). To provide greater flexibility over the search output, the advanced search can be used to filter the search output based on multiple search queries, such as the lncRNA and cancer name, lncRNA expression pattern, and the number of transcript variants of lncRNA (Figure S1 A, D). In addition to using the cancer name in the general search bar and advanced search to search the database, the tissue-based cancer distribution in the interactive human anatomy can be used to generate similar search output with respect to the cancer of the selected tissue/ organ (Figure S1 E).

#### B. LncRNA G4 prediction

The database offers a comprehensive G4 prediction for each catalogued lncRNA and their transcript variants, employing well-established G4-prediction tools such as QGRS mapper and G4Hunter (Figure 1A). The database search output fetches the details of searched lncRNA or the lncRNAs associated with the searched cancer, with regards to their transcript variants and corresponding NCBI accession IDs obtained from NCBI Nucleotide (Figure S1 B, C, F).^74^ The NCBI accession IDs can also be used to navigate to the source details of the FASTA sequence (Figure S1 F). Thereafter, using either or both G4-prediction tools, the output furnishes an exhaustive list of all possible PQS within these lncRNAs and their transcript variants. The output also presents the position, length, and G4 propensity score of listed PQS. Additionally, the database categorizes the PQS predicted by G4-prediction tools into anticipated G4 types (2G, 3G, and 4G), providing structural insights into the probable G4 motifs (Figure S1 G, H). The database also facilitates the comparison of PQS predicted by the two G4-prediction tools and enables the re-analysis of catalogued lncRNA sequences by modifying the analysis parameters of both G4-prediction tools (Figure S1 I). Furthermore, the G4-prediction tools are integrated as standalone tools, empowering users to critically assess, categorize, and compare G4 predictions for any given sequence and NCBI accession ID.

#### C. LncRNA Subcellular Localization

CanLncG4 also provides valuable insights into the subcellular localization of catalogued lncRNAs across diverse cell lines. The database search output provides cytoplasmic to nuclear localization: relative concentration index, expression values, and distribution data, sourced from the LncATLAS database, for the searched lncRNA or the lncRNAs associated with the searched cancer (Figure S1 B, C, J, K).^75^

#### D. Statistics, downloads, glossary, and help section

Furthermore, the detailed statistics of the catalogued lncRNA-cancer data, and the meta-analysis of G4-prediction, can be visualized from the statistics section (Figure S2 A-E). These statistics offer valuable insights into the G4 landscape within lncRNAs across various human cancers. Additionally, they provide an overview of the effectiveness of the parameters employed in G4-prediction tools for identifying PQS. The glossary section provides the list of all cancers and its subtype, and the lncRNAs catalogued in the database, which can be selected to direct to their respective search outputs (Figure S3 A, B). All the raw and meta-analyzed data can be downloaded from the download section of the database (Figure S3 C). The help section can be referred to obtain details concerning the website navigation and usage (Figure S3 D).

### Workflow for identification of LncRNA-G4 interacting Partners

CanLncG4 conducts a comprehensive meta-analysis of experimentally validated interacting partners, both RNA and protein, associated with the catalogued lncRNAs, relying on the recently available data from NPInter v4.0 and LncTarD 2.0.^71, 72^ Additionally, the database imparts information about the experimentally validated RNA G4-binding proteins (RGBP) interacting with the catalogued lncRNAs, utilizing QUADRatlas, G4IPDB, and scientific literature mining (Figure 1B).^64, 69, 76^ The search bar in the lncRNA-G4 interacting partner section of the database can be used to search the database by the lncRNA name (Figure S4 A). The search output presents the details of the established lncRNA-protein and lncRNA-RNA interactions, and provides an option to obtain the details of the RNA G4-binding capabilities of the protein interacting with the searched lncRNA (Figure S4 B-F). Since a majority of the lncRNAs catalogued in the database contain putative G4-forming regions, detail of proteins interacting with these putative G4-forming lncRNAs in conjunction with the established RNA G4-binding potential of such proteins can be used as a lead to investigate the G4 mediated lncRNA-protein interaction.

### Human cancer-dysregulated Putative Quadruplex-forming Sequence harboring lncRNAs forms parallel and stable G4s *in vitro*

We validated our *in silico* G4-prediction pipeline by performing *in vitro* studies on high-scoring PQS from randomly selected *HOXA-AS2*, *DLX6-AS1*, *GHET1*, *LINP1*, and *LINC00504* lncRNAs that are dysregulated in multiple human cancers (Figure 2A, B, Table S1). We have referred to these PQSs hereafter by the name of their respective lncRNAs. We prepared the *HOXA-AS2*, *DLX6-AS1*, *GHET1*, *LINP1*, and *LINC00504* lncRNAs by *in vitro* transcription (IVT) as described in the Experimental section. The design of template DNA oligonucleotides used in this work mirrors the design of templates that has been used in our recently communicated works. The sequences of cDNA antisense templates for *HOXA-AS2*, *DLX6-AS1*, *GHET1*, *LINP1*, and *LINC00504* lncRNA PQS used in this validation exercise are listed in Table S1.

**Figure 2.**
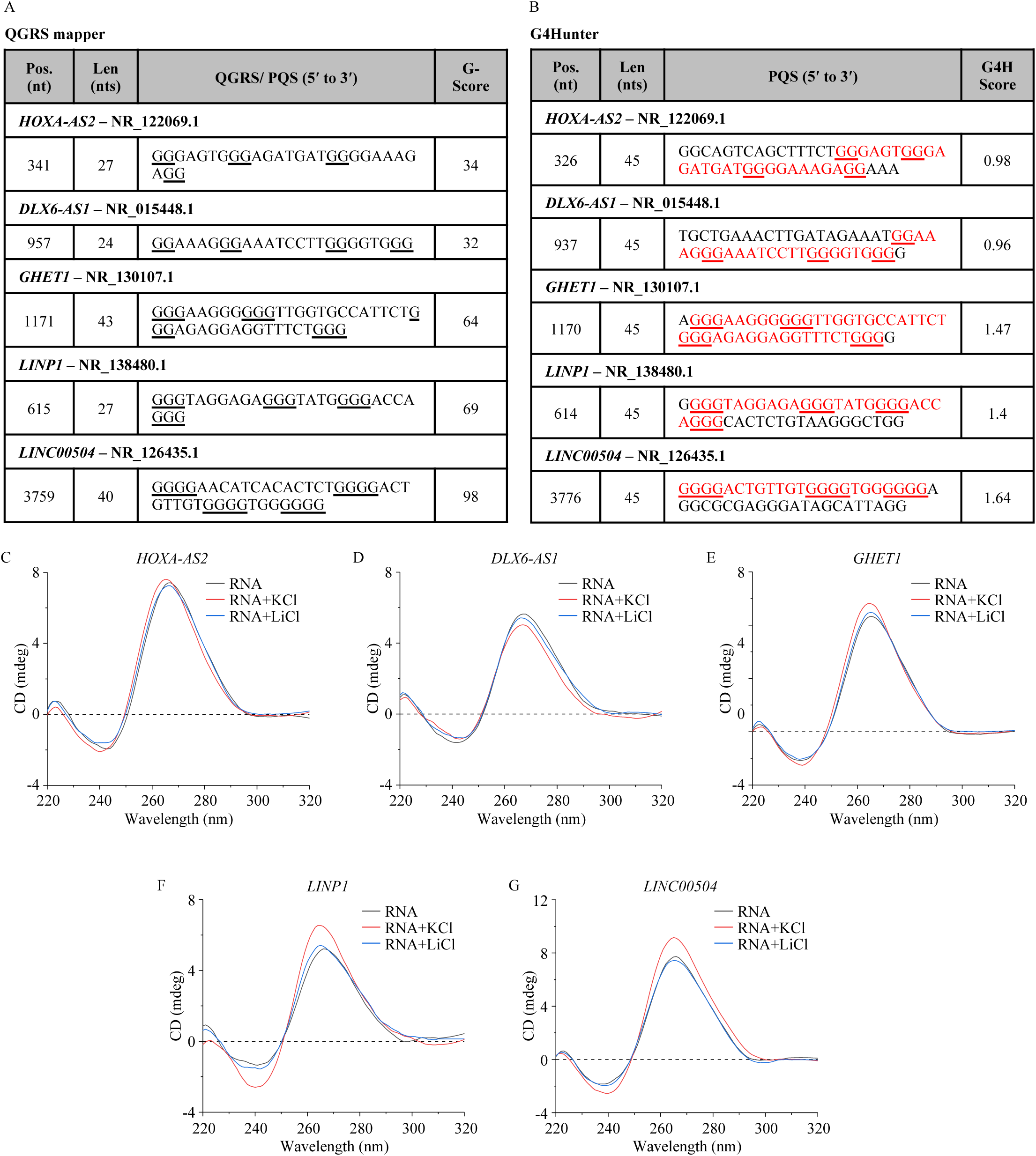
*In vitro* formation of parallel G4s in *HOXA-AS2*, *DLX6-AS1*, *GHET1*, *LINP1*, and *LINC00504* lncRNAs. A-B) CanLncG4 identified PQS from selected lncRNAs by employing A) QGRS mapper, and B) G4Hunter. C-G) CD spectroscopy of IVT RNAs (5 µM) folded in the presence or absence of 100 mM KCl or LiCl. Maxima and minima in mean CD spectra at ca. 265 and 240 nm, respectively, correspond to parallel G4 topologies.

CD spectroscopy has emerged as a widely employed technique for characterizing the formation and unravelling the topology of G-quadruplexes (G4s) in both DNA and RNA molecules.^77^ CD spectra of all the folded IVT RNAs (5 µM) displayed maxima and minima at ca. 265 and 240 nm, suggesting the formation of parallel G4 structures in each case.^77^ The parallel G4 topology of *HOXA-AS2*, *DLX6-AS1*, *GHET1*, *LINP1*, and *LINC00504* lncRNAs (Figure 2C-G) is aligned with the most commonly formed topology of RNA G4s.^28^ We prepared the folded RNA samples with or without monovalent ions (100 mM of K^+^ and Li^+^) to assess their effect on G4 stability.^78, 79^ In the presence of KCl, the CD spectra of *HOXA-AS2*, *GHET1*, *LINP1*, and *LINC00504* lncRNAs displayed a modest increase in the intensity of CD band at 265 nm (Figure 2C, E-G). In contrast, *DLX6-AS1* lncRNA displayed a weaker maxima at 265 nm in the presence of K^+^ (Figure 2D). The CD maxima at 265 nm for *HOXA-AS2*, *DLX6-AS1*, and *LINC00504* lncRNAs was lowered in the presence of LiCl in the folding buffer, when compared to the absence of additional monovalent cations (Figure 2C, D, G). Interestingly, *GHET1* and *LINP1* displayed a modest increase in the intensity of the CD band at 265 nm in the presence of Li^+^ (Figure 2E, F). While Li^+^ is conventionally associated with the destabilization of G4s, alternative effects have been reported previously.^78–80^ These results are also aligned with our previously communicated results pertaining to the behaviour of PQS of dysregulated lncRNAs in CRC. Overall, all five randomly selected lncRNAs are capable of folding into parallel G4s albeit with variable effects of monovalent cations on these structures.

The porphyrin TMPyP4, a well-established DNA-G4 ligand, is recognized for its ability to destabilize G4 structures across various classes of RNA.^81–84^ We investigated the effect of TMPyP4 on the G4s formed by *HOXA-AS2*, *DLX6-AS1, GHET1, LINP1,* and *LINC00504* lncRNAs. The folded IVT RNAs (5 µM) were titrated with TMPyP4 till a final concentration of 25 µM. CD spectra were measured on the samples after each titration. All the RNAs, namely *HOXA-AS2*, *DLX6-AS1*, *GHET1*, *LINP1*, and *LINC00504* lncRNAs displayed a progressive reduction in CD band intensities upon titration with TMPyP4 (Figure 3A-E). This behaviour is in agreement with the reported behaviour of RNA G4s with TMPyPy.^82, 83^ Among these lncRNAs, *DLX6-AS1* displayed a significant decrease in CD band intensity upon titration with TMPyP4 (Figure 3B). Nevertheless, none of the lncRNAs displayed complete flattening of their CD spectra even in the presence of 25 µM of TMPyP4 (Figure 3A-E). The persistence of CD bands of *HOXA-AS2*, *GHET1*, *LINP1*, *LINC00504* lncRNAs, and to a limited extent of *DLX6-AS1* lncRNA indicate their ability to maintain parallel G4 topology even at high concentrations of TMPyP4.

**Figure 3.**
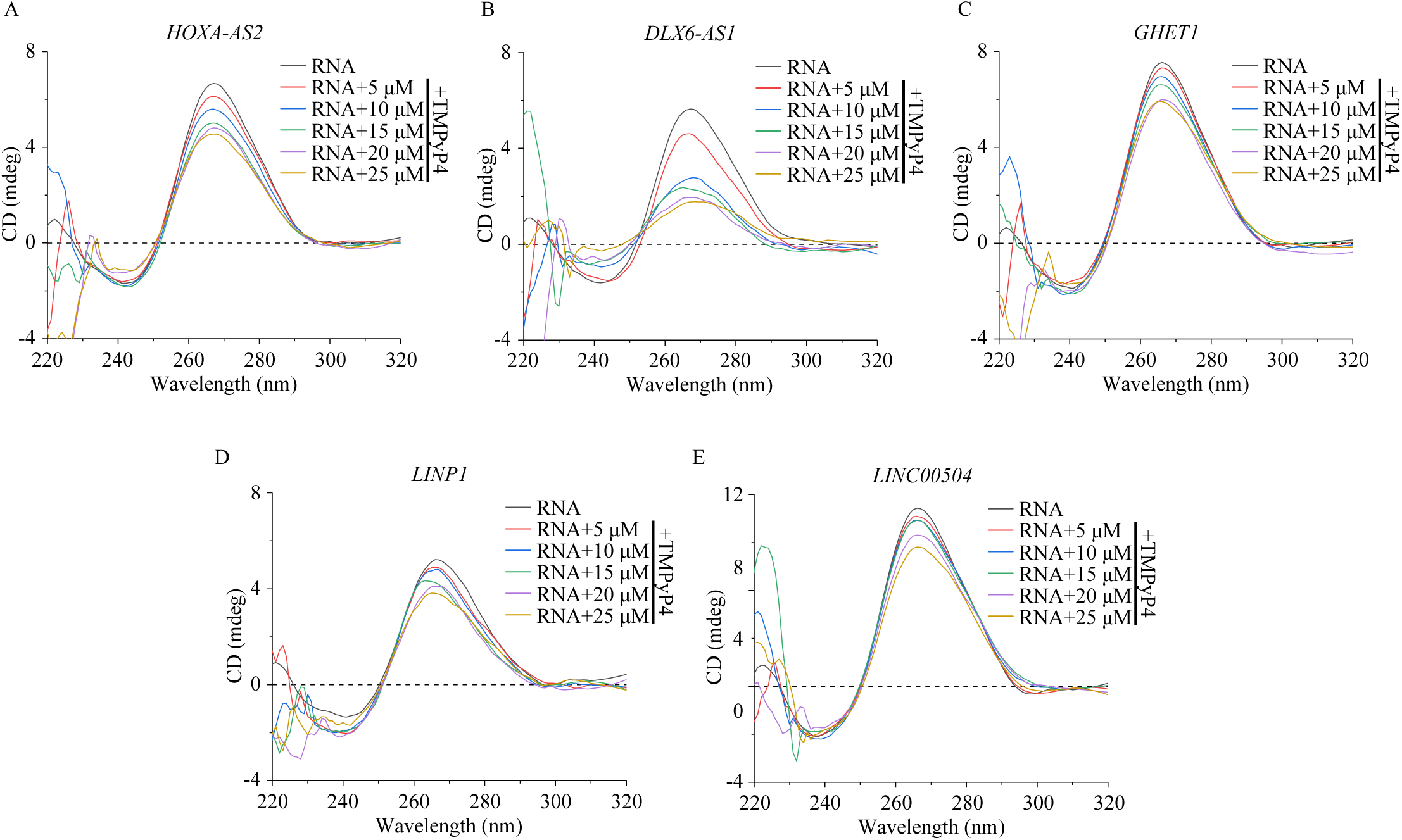
*In vitro* formation of stable G4s in *HOXA-AS2*, *DLX6-AS1*, *GHET1*, *LINP1*, and *LINC00504* lncRNAs. A-E) CD spectroscopy of folded IVT RNAs (5 µM) titrated with TMPyP4 (5 – 25 µM). No flattening of mean CD spectra shows the formation of stable parallel G4 topologies.

We used the ThT fluorescence enhancement assay to assess the effect of monovalent cations on the stability of G4s formed by *HOXA-AS2*, *DLX6-AS1*, *GHET1*, *LINP1*, and *LINC00504* lncRNAs. Thioflavin T (ThT) has been effectively utilized to detect RNA G4s due to its capacity to stack onto the terminal G-quartets, resulting in a notable increase in fluorescence signal compared to other structural conformations of RNA.^85^ The ThT assay was performed by adding ThT (2 µM) to IVT RNAs (2 µM) suitably folded in the presence or absence of 100 mM KCl/LiCl, followed by fluorescence excitation and emission measurements. The fold of enhancement of ThT upon interaction with the G4s in *HOXA-AS2*, *DLX6-AS1*, *GHET1*, *LINP1*, and *LINC00504* lncRNAs in comparison to ThT alone were estimated to be 221, 218, 289, 218, and 180-fold, respectively (Figure 4A-F). These handsome ThT fluorescence enhancements validate G4 formation in each case.

**Figure 4.**
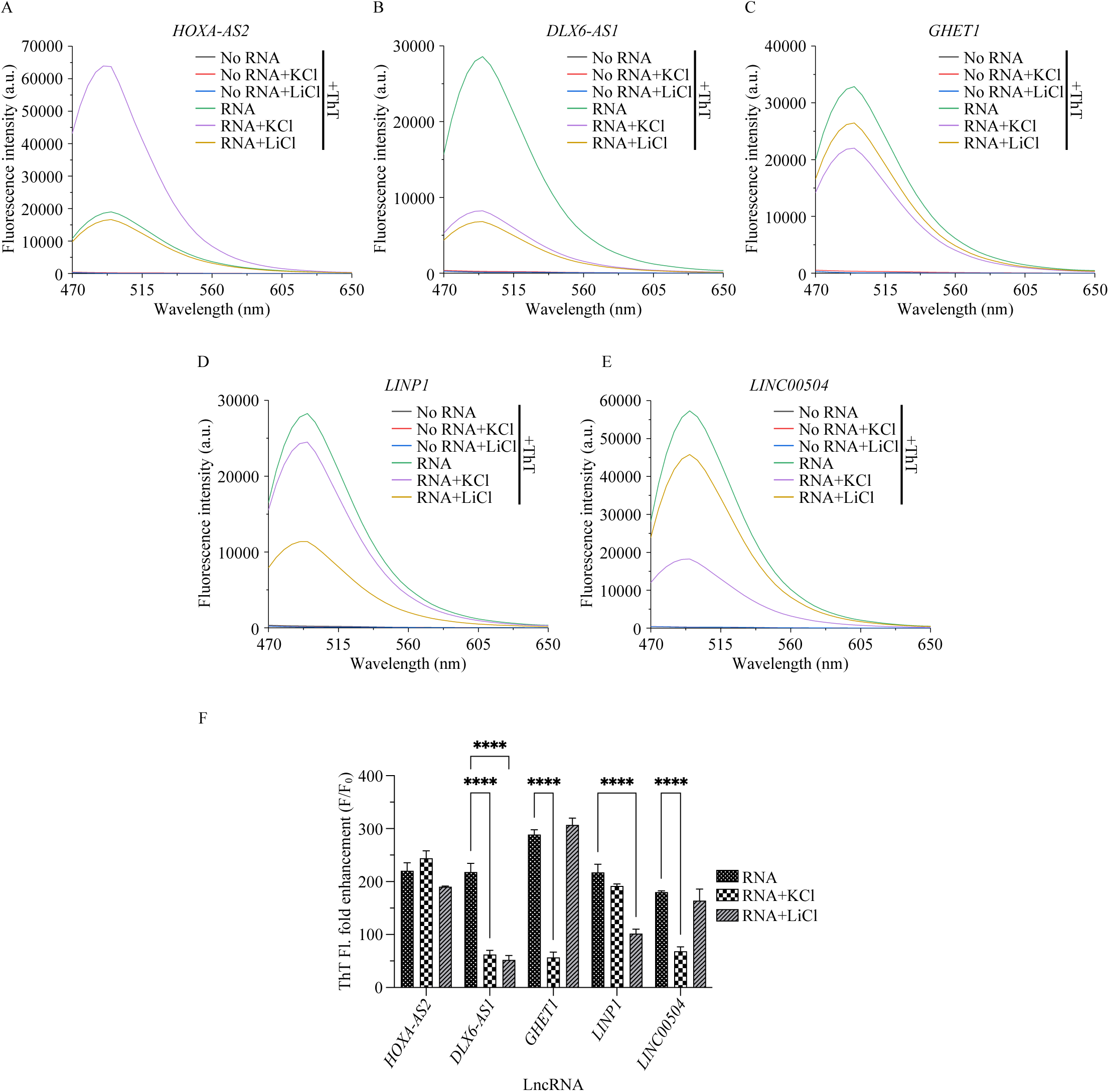
Monovalent cations affect the stability of *HOXA-AS2*, *DLX6-AS1*, *GHET1*, *LINP1*, and *LINC00504* lncRNA G4s. A-F) ThT fluorescence enhancement assay of IVT RNAs (2 µM) folded in the presence or absence of 100 mM KCl or LiCl with ThT (2 µM). A-E) Increased mean fluorescence emission spectra, and F) fold enhancement in mean ± SEM ThT fluorescence (F/ F0: ThT fluorescence in the presence of RNA/ absence of RNA) at 488 nm when excited at 445 nm, corresponds to G4 formation in the presence or absence of monovalent cations. P-values: 0.0332 (*), 0.0021 (**), 0.0002 (***), <0.0001 (****). Non-significant P-values are not represented.

KCl and LiCl exerted interesting effects on the G4 stability of the RNAs as evident from the corresponding fold of ThT fluorescence enhancement (Figure 4A-F). Apart from a greater variation in the fold of ThT enhancement across the lncRNAs, the magnitude of enhancements observed in the presence of KCl did not follow the same register as in the absence of ions. For example, while *HOXA-AS2* displayed a 244-fold enhancement of ThT fluorescence, *DLX6-AS1* only displayed a 62-fold enhancement. The corresponding magnitudes in the absence of KCl were 221 and 218, for *HOXA-AS2* and *DLX6-AS1*, respectively. *HOXA-AS2* lncRNA manifests a superior stabilizing effect of K^+^ compared to the other lncRNAs. Li^+^ exerted an intriguing stabilizing effect on *GHET1* based on the observed 307-fold enhancement of ThT fluorescence. Notably, the other lncRNAs displayed a lower fold of ThT fluorescence enhancement in the presence of Li^+^. Notwithstanding the canonical G4-destabilizing behavior of Li^+^ towards *GHET1*, the other G4s clearly possess suitable topologies for coordination by Li^+^. Such unexpected action of Li^+^ has been reported previously, including in our recent communication on G4s in dysregulated lncRNAs of CRC.^78–80^

## Conclusion

The dysregulation of lncRNAs has been strongly implicated in various diseases, particularly cancer.^45, 48^ G4 harboring lncRNAs have emerged as important participants and potentially significant contributors in cancer development.^41, 48–50^ Recent studies have also emphasized the importance of lncRNA G4-protein interactions in regulating gene expression and promoting cancer progression.^25, 40, 42, 45^ Our current work is motivated by the need for exploring G4-formation by lncRNAs and the identification of potential roles in the context of cancer.

Advancements in genome sequencing technologies have facilitated the development of sequence-based computational prediction tools for identifying putative G4 motifs in DNA and RNA.^51^ These existing tools are based on regular expression matching, scoring and ranking sequences, and machine learning methods.^51–62^ Considering the diversity of informatics resources capturing lncRNA-cancer associations and computational tools for G4-prediction and assessment, we sought to create a platform that would integrate diverse data types and ultimately shed light on the G4-forming potential of lncRNAs dysregulated in cancers.^63–72^

In this work, we introduce CanLncG4, a platform that consolidates experimentally validated associations between lncRNAs and various human cancers. CanLncG4 offers extensive G4-prediction analysis for each lncRNA transcript variant, categorizing predicted G4 motifs and providing valuable structural insights. Additionally, CanLncG4 provides insights into lncRNA subcellular localization, a meta-analysis of RNAs and proteins interacting with lncRNAs, and information on the RNA G4-binding capabilities of these proteins. We also validated our *in silico* G4-prediction pipeline by performing *in vitro* studies on high-scoring PQS from randomly selected cancer-dysregulated lncRNAs. The finding suggests the formation of G4s in these lncRNAs and, consequently, displays the G4-prediction potential of our methodology. CanLncG4 provides a valuable platform for conducting preliminary screening of cancer-associated lncRNAs, particularly focusing on their potential to form G4s and their interacting partners. By leveraging this platform, researchers can efficiently identify and prioritize lncRNA candidates that exhibit promising characteristics for further *in vitro* and *in cellulo* investigations. This streamlined approach accelerates the process of discovering lncRNAs with diagnostic and therapeutic potential in cancer research. Looking ahead, the continual expansion of CanLncG4 to include lncRNAs associated with additional cancers will broaden its applicability and contribute to advancing our understanding of the intricate roles played by lncRNAs in cancer biology.

Together, CanLncG4 can serve as a comprehensive platform for accessing information about cancer-dysregulated lncRNAs, their G4-forming potential and interacting partners and, hence, would likely aid in substantive understanding on the question of - <u>Can</u>cer-<u>Lnc</u>RNA form <u>G4</u>?

## Data Availability

The data is available at https://canlncg4.com/.

## Acknowledgements

The authors thank Department of Biological Sciences and Engineering, Department of Chemistry, Common Research & Technology Development Hub, and Central Instrumentation facility at Indian Institute of Technology Gandhinagar, Gandhinagar, Gujarat, India for providing instrumentation facility. The authors also thank Angshuman Mandal for technical assistance with data collection concerning the LncRNA-G4 Interacting Partners. All the illustrations are Created with BioRender.com.

