## Supplemental Figures for "CanLncG4: A database curated for the assessment of G4s in the lncRNAs dysregulated in various human cancers"

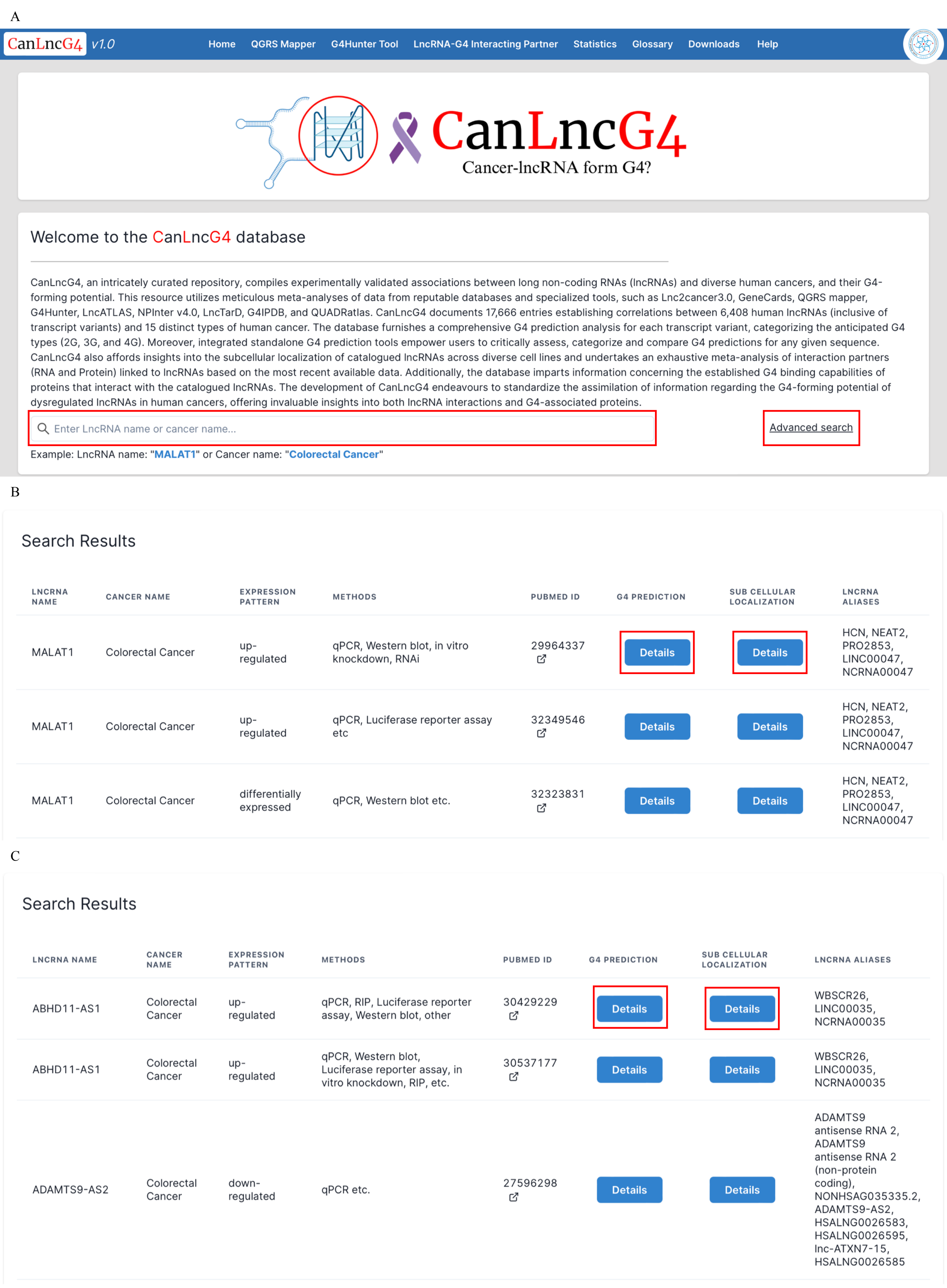

A
CanLncG4 v1.0
Home
QGRS Mapper
G4Hunter Tool
LncRNA-G4 Interacting Partner
Statistics
Glossary
Downloads
Help
CanLncG4
Cancer-lncRNA form G4?
Welcome to the CanLncG4 database
CanLncG4, an intricately curated repository, compiles experimentally validated associations between long non-coding RNAs (lncRNAs) and diverse human cancers, and their G4-forming potential. This resource utilizes meticulous meta-analyses of data from reputable databases and specialized tools, such as Lnc2cancer3.0, GeneCards, QGRS mapper, G4Hunter, LncATLAS, NPInter v4.0, LncTarD, G4IPDB, and QUADRatlas. CanLncG4 documents 17,666 entries establishing correlations between 6,408 human lncRNAs (inclusive of transcript variants) and 15 distinct types of human cancer. The database furnishes a comprehensive G4 prediction analysis for each transcript variant, categorizing the anticipated G4 types (2G, 3G, and 4G). Moreover, integrated standalone G4 prediction tools empower users to critically assess, categorize and compare G4 predictions for any given sequence. CanLncG4 also affords insights into the subcellular localization of catalogued lncRNAs across diverse cell lines and undertakes an exhaustive meta-analysis of interaction partners (RNA and Protein) linked to lncRNAs based on the most recent available data. Additionally, the database imparts information concerning the established G4 binding capabilities of proteins that interact with the catalogued lncRNAs. The development of CanLncG4 endeavours to standardize the assimilation of information regarding the G4-forming potential of dysregulated lncRNAs in human cancers, offering invaluable insights into both lncRNA interactions and G4-associated proteins.
Enter LncRNA name or cancer name...
Advanced search
Example: LncRNA name: "MALAT1" or Cancer name: "Colorectal Cancer"
B
Search Results
LNCRNA NAME
CANCER NAME
EXPRESSION PATTERN
METHODS
PUBMED ID
G4 PREDICTION
SUB CELLULAR LOCALIZATION
LNCRNA ALIASES
MALAT1
Colorectal Cancer
up-regulated
qPCR, Western blot, in vitro knockdown, RNAi
29964337
Details
Details
HCN, NEAT2, PRO2853, LINC00047, NCRNA00047
MALAT1
Colorectal Cancer
up-regulated
qPCR, Luciferase reporter assay etc
32349546
Details
Details
HCN, NEAT2, PRO2853, LINC00047, NCRNA00047
MALAT1
Colorectal Cancer
differentially expressed
qPCR, Western blot etc.
32323831
Details
Details
HCN, NEAT2, PRO2853, LINC00047, NCRNA00047
C
Search Results
LNCRNA NAME
CANCER NAME
EXPRESSION PATTERN
METHODS
PUBMED ID
G4 PREDICTION
SUB CELLULAR LOCALIZATION
LNCRNA ALIASES
ABHD11-AS1
Colorectal Cancer
up-regulated
qPCR, RIP, Luciferase reporter assay, Western blot, other
30429229
Details
Details
WBSCR26, LINC00035, NCRNA00035
ABHD11-AS1
Colorectal Cancer
up-regulated
qPCR, Western blot, Luciferase reporter assay, in vitro knockdown, RIP, etc.
30537177
Details
Details
WBSCR26, LINC00035, NCRNA00035
ADAMTS9-AS2
Colorectal Cancer
down-regulated
qPCR etc.
27596298
Details
Details
ADAMTS9 antisense RNA 2, ADAMTS9 antisense RNA 2 (non-protein coding), NONHSAG035335.2, ADAMTS9-AS2, HSALNG0026583, HSALNG0026595, lnc-ATXN7-15, HSALNG0026585

D

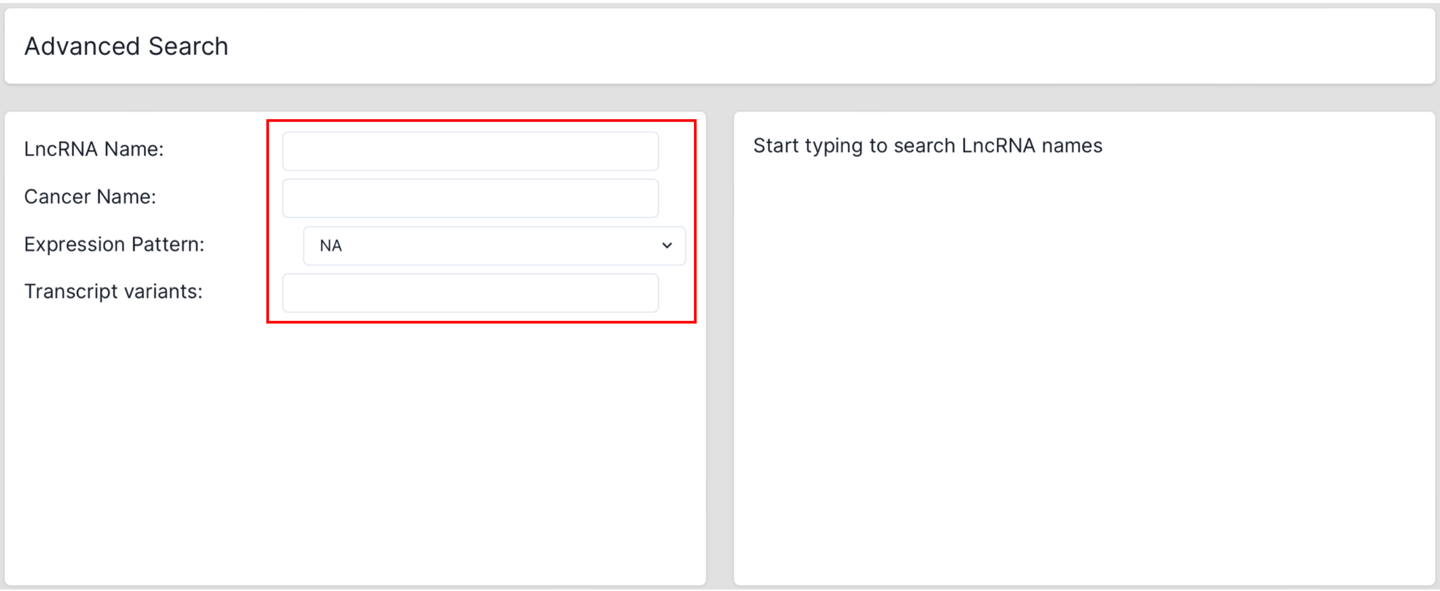


E

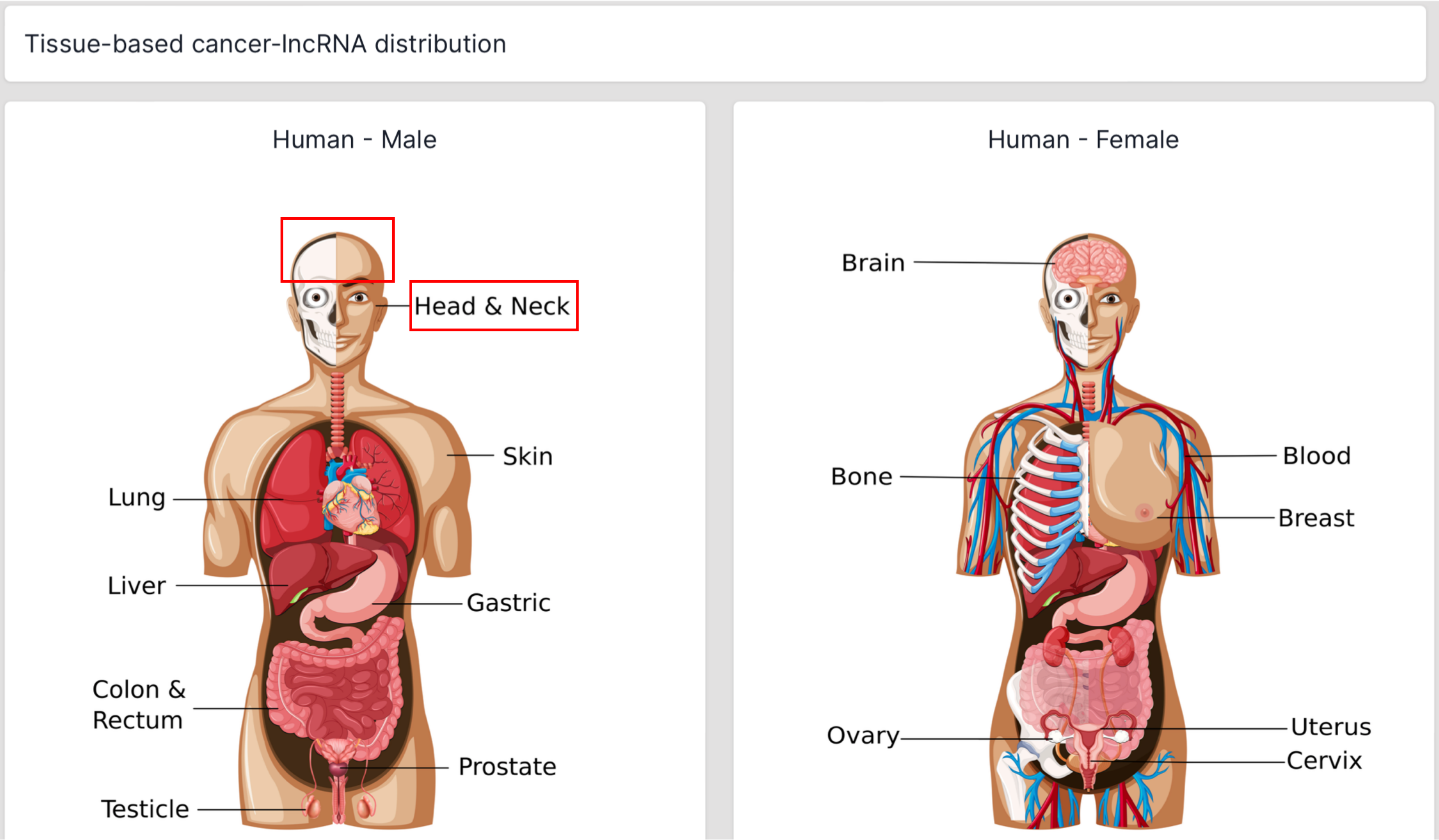


F

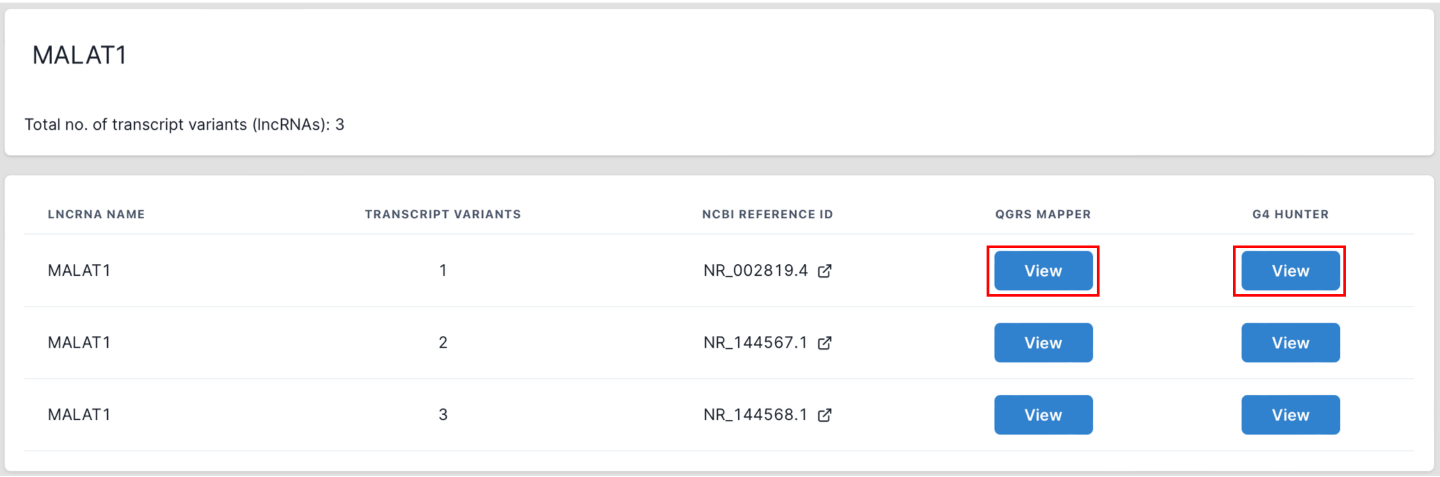


| LNCRNA NAME | TRANSCRIPT VARIANTS | NCBI REFERENCE ID | QGRS MAPPER | G4 HUNTER |
|---|---|---|---|---|
| MALAT1 | 1 | NR_002819.4 | View | View |
| MALAT1 | 2 | NR_144567.1 | View | View |
| MALAT1 | 3 | NR_144568.1 | View | View |

G

Max Length: - 45 +

Min G-group: 2

Loop size: 0 to 36

Analyze

| TOTAL NO. OF PQS | NO. OF 2G PQS | NO. OF 3G PQS | NO. OF 4G PQS |
|---|---|---|---|
| 26.5 | 23 | 3.5 | 0 |

| POSITION | LENGTH | TYPE OF G-QUADRAPLEX | G-SCORE | SEQUENCE |
|---|---|---|---|---|
| 30 | 21 | 2G | 26 | GGACTGGGGCCCCGCAACTGG |
| 128 | 34 | 2G | 21 | GGCAGGTCCCCTCTGACGCCTCCGGGAGCCCAGG |
| 221 | 22 | 2G | 30 | GGCTGGCCATTCCAGGTGGTGG |

H

Window size: - 45 +

Threshold: 0.9

Analyze

| TOTAL NO. OF PQS | NO. OF 2G PQS | NO. OF 3G PQS | NO. OF 4G PQS |
|---|---|---|---|
| 13 | 18 | 4 | 0 |

| POSITION | LENGTH | TYPE OF G-QUADRAPLEX | G-SCORE | SEQUENCE |
|---|---|---|---|---|
| 589 | 45 | 3G | 1.24 | GGGCCGTGGGGGGCTGGCGGCAACTGGGGGGCCGCAGATCAGAGT |
| 659 | 45 | 2G | 1.16 | GGGGCTCAGGCGGGGAGCAGCTCTGTGGTGTGGGATTGAGGCGTT |
| 1303 | 45 | 2G | 0.93 | CGGTAGGCATTGAGGCAGCCAGCGCAGGGGCTTCTGCTGAGGGGG |

I

Max Length: - 45 +

Min G-group: 2

Loop size: 0 to 36

Analyze

| TOTAL NO. OF PQS | NO. OF 2G PQS | NO. OF 3G PQS | NO. OF 4G PQS |
|---|---|---|---|
| 39.5 | 41 | 7.5 | 0 |

| POSITION | LENGTH | TYPE OF G-QUADRAPLEX | G-SCORE | SEQUENCE |
|---|---|---|---|---|
| 30 | 21 | 2G | 26 | GGACTGG |
| 128 | 34 | 2G | 21 | GGCAGGT |
| 221 | 22 | 2G | 30 | GGCTGGC |
| 282 | 12 | 2G | 33 | GGGGTTT |
| 421 | 42 | 2G | 31 | GGTGTGT |
| 590 | 31 | 3G | 60 | GGGCCGT |
| 660 | 41 | 2G | 36 | GGGGCTC |

Window size: - 45 +

Threshold: 0.9

Analyze

| TOTAL NO. OF PQS | NO. OF 2G PQS | NO. OF 3G PQS | NO. OF 4G PQS |
|---|---|---|---|
| 13 | 18 | 4 | 0 |

| POSITION | LENGTH | TYPE OF G-QUADRAPLEX | G-SCORE | SEQUENCE |
|---|---|---|---|---|
| 589 | 45 | 3G | 1.24 | GGGCCGT |
| 659 | 45 | 2G | 1.16 | GGGGCTC |
| 1303 | 45 | 2G | 0.93 | CGGTAGG |
| 1312 | 45 | 2G | 0.96 | TTGAGGC |
| 1322 | 45 | 2G | 0.98 | CAGCGCA |
| 3102 | 45 | 2G | 1.13 | TAATGGG |
| 3432 | 45 | 2G | 0.93 | GTTTTGTA |

J

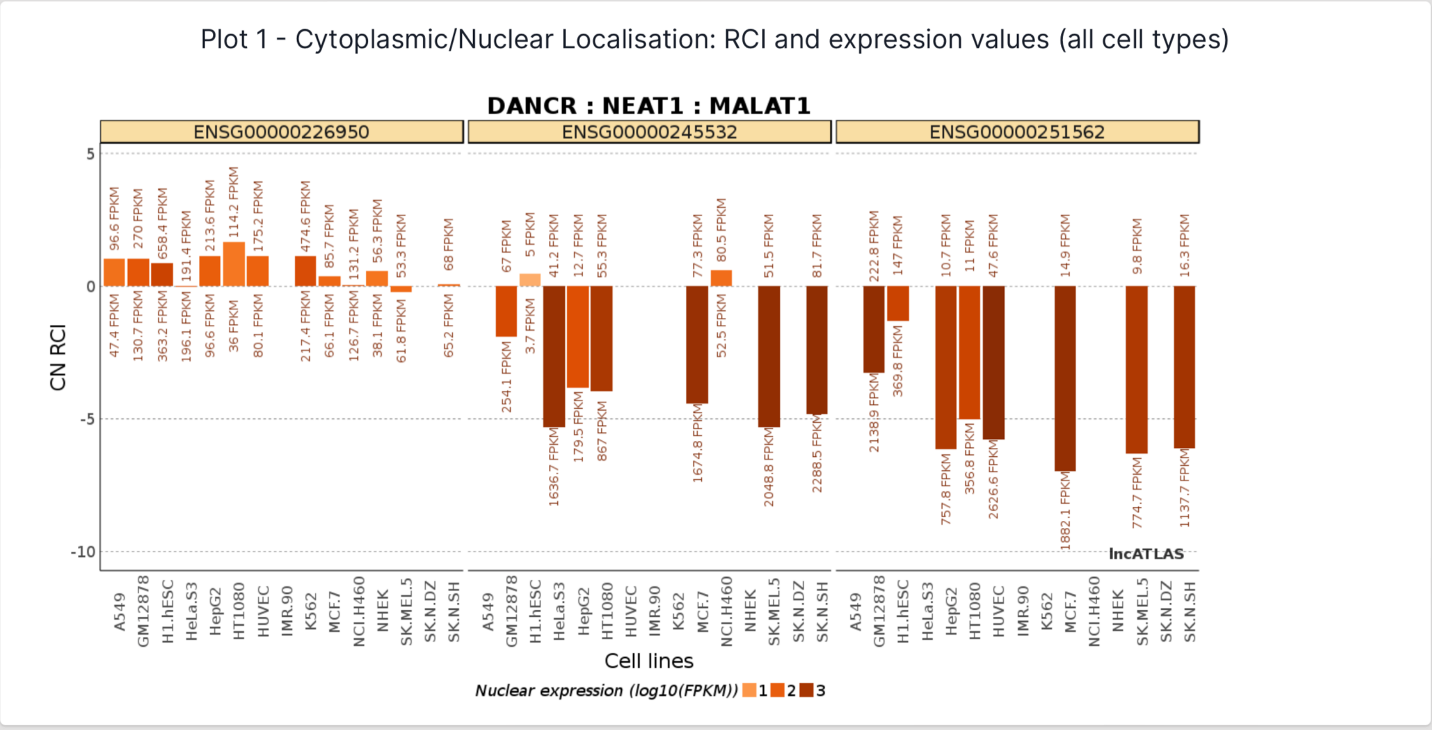


K

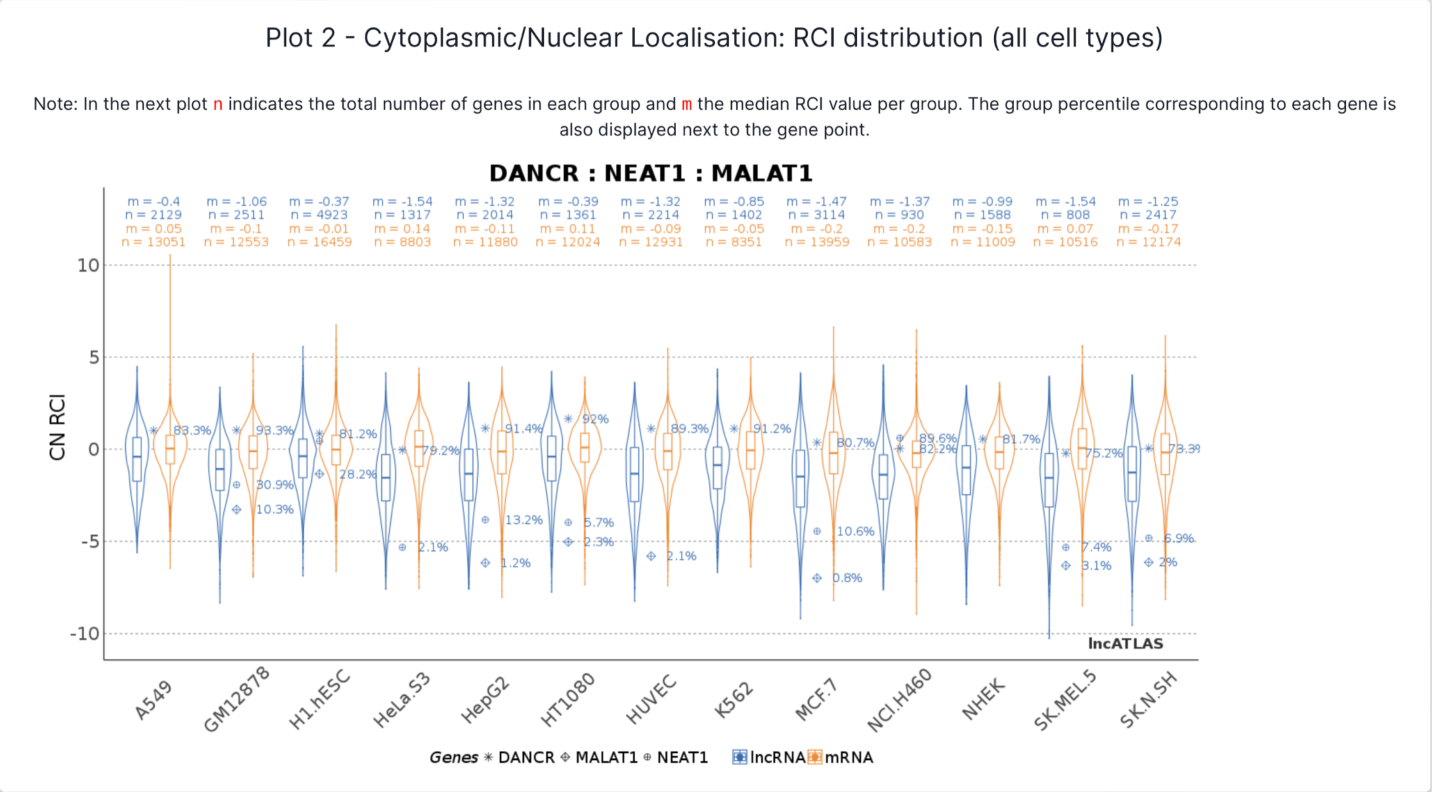


**Figure S1. LncRNA-cancer association, G4-prediction, and subcellular localization sections of CanLncG4.** A) Homepage with a general search bar and advanced search. B-C) Search output with A) lncRNA name, C) cancer name as a query. D) Parameters of advanced search. E) Interactive SVGs to obtain tissue/ organ-based cancer-specific search output. F) Details of lncRNA transcript variants and their NCBI accession IDs. G-I) G4-prediction of searched lncRNA using G) QGRS mapper, H) G4Hunter, I) comparison of PQS obtained from QGRS mapper and G4Hunter. J-K) Cytoplasmic to nuclear localization plots from LncATLAS. The red box highlights the tabs that can be used to proceed to the next section.

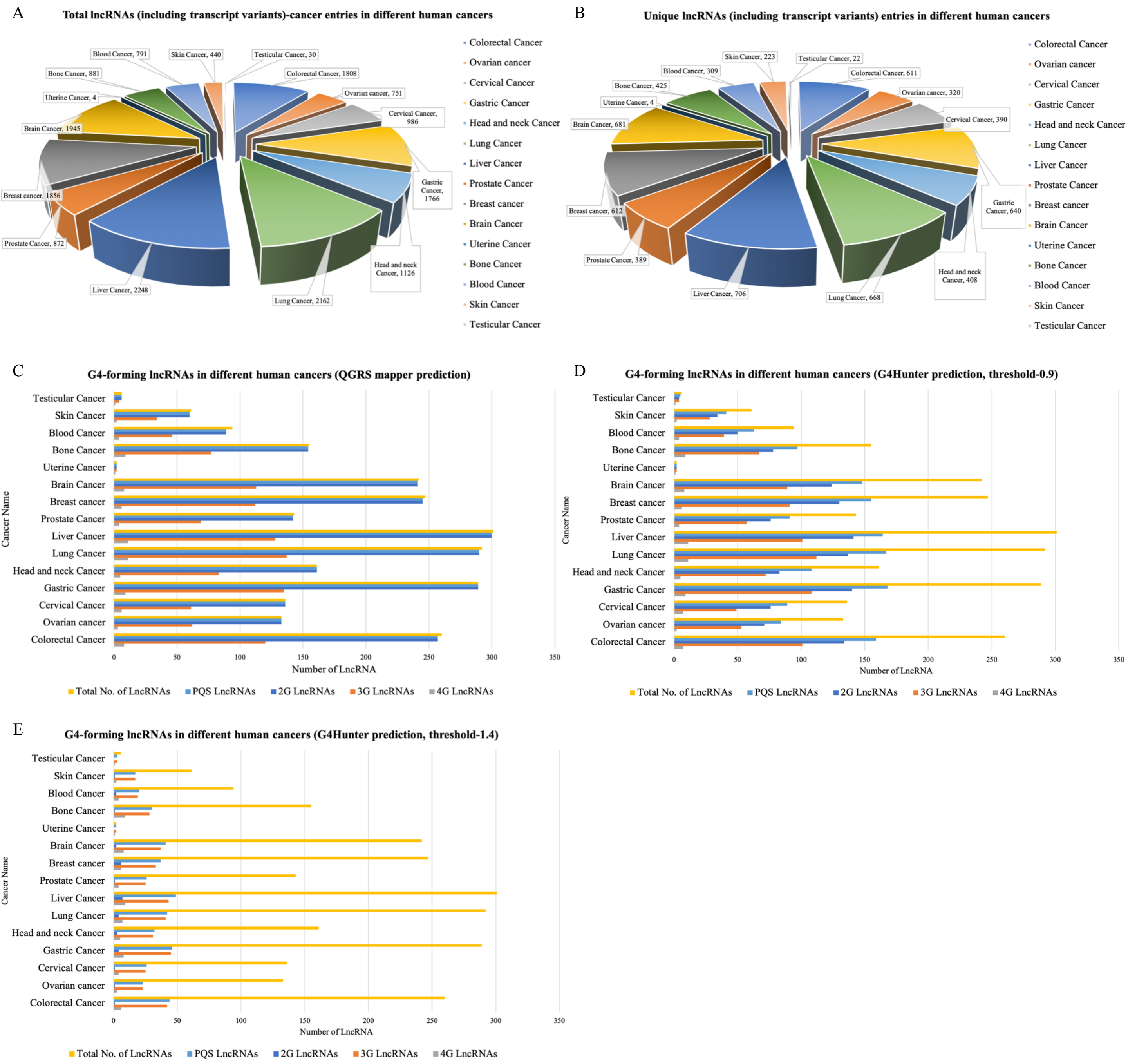


**Figure S2. Statistics of data catalogued and meta-analyzed in CanLncG4.** A-B) LncRNA-cancer associations. C-E) G4-prediction of lncRNAs using C) QGRS mapper, D-E) G4Hunter.

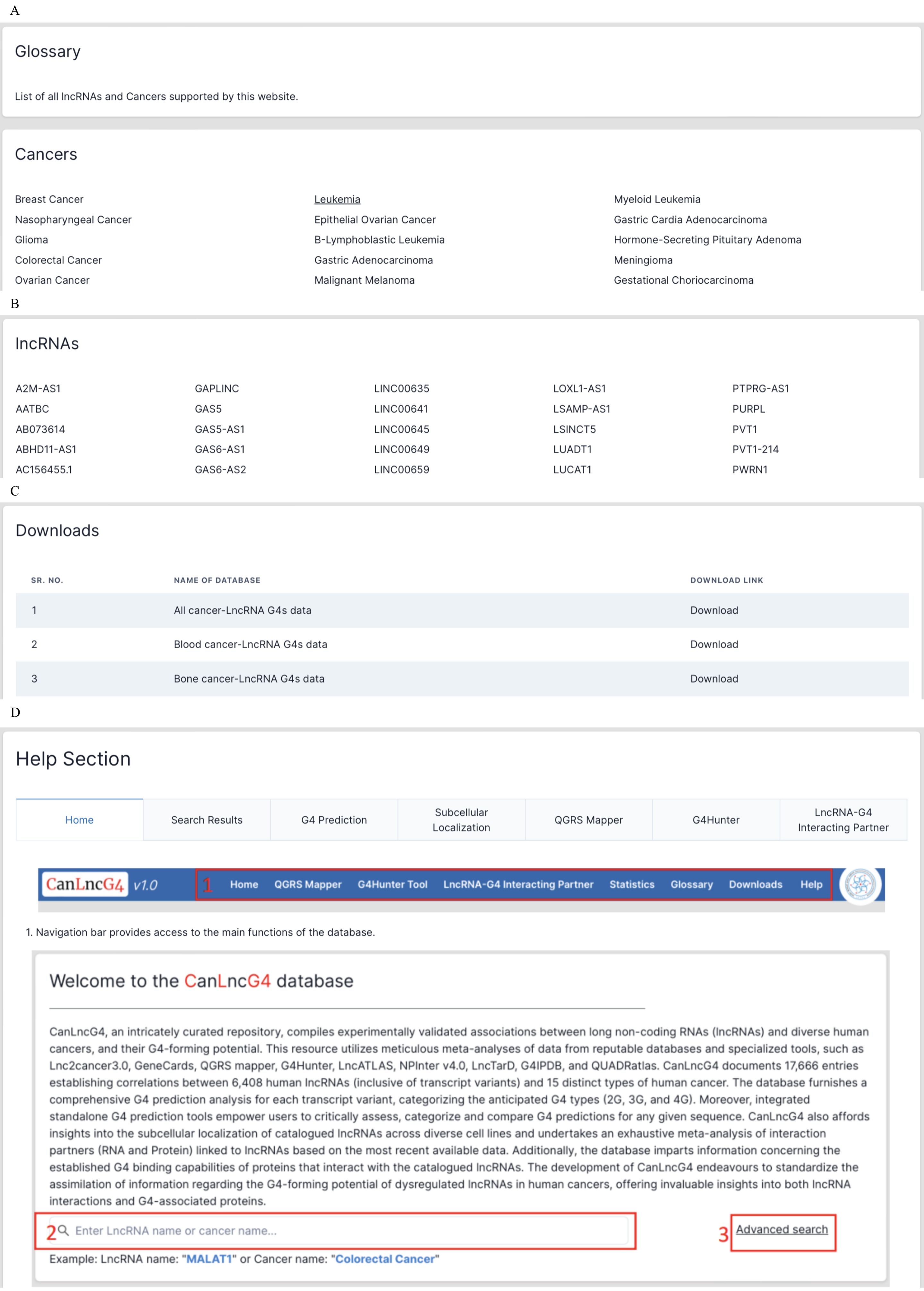


**Figure S3. Other sections of CanLncG4.** A-B) Glossary for A) cancer names and their sub-types, B) lncRNA names. C) Downloads. D) Help.

A

LncRNA - G4 Interacting Partner

Enter LncRNA

Search

B

LncRNA-protein interactions - NPInter

| NPINTER INTERACTION ID | INTERACTOR NAME | INTERACTOR TYPE | INTERACTOR ID | TARGET NAME | TARGET TYPE | KNOWN G4 BINDER? | TARGET ID | INTERACTION MECHANISM | INTERACTION LEVEL | INTERACTION CLASS |
|---|---|---|---|---|---|---|---|---|---|---|
| ncRI-40004559 | MALAT1 | lncRNA | NONHSAG008675 | ELAVL1 | protein | View Details | P70372 | ncRNA-protein binding | RNA-Protein | binding |
| ncRI-40007018 | MALAT1 | lncRNA | NONHSAG008675 | CASP8 | protein | View Details | Q14790 | ncRNA-protein binding | RNA-Protein | binding |
| ncRI-40007019 | MALAT1 | lncRNA | NONHSAG008675 | CASP3 | protein | View Details | P42574 | ncRNA-protein binding | RNA-Protein | binding |

C

LncRNA-Protein Interactions - LncTarD

| REGULATION ID | REGULATOR NAME | REGULATOR TYPE | REGULATOR ENSEMBLE ID | REGULATOR ALIASES | TARGET NAME | TARGET TYPE | TARGET ENSEMBLE ID | TARGET ALIASES | KNOWN G4 BINDER? | REGULATORY MECHANISM |
|---|---|---|---|---|---|---|---|---|---|---|
| RID02608 | MALAT1 | lncRNA | ENSG00000251562 | HCN\|LINC00047\|NCRNA00047\|NEAT2\|PRO2853 | ELAVL1 | PCG | ENSG00000066044 | ELAV1\|HUR\|Hua\|MelG | View Details | interact with protein |
| RID02609 | MALAT1 | lncRNA | ENSG00000251562 | HCN\|LINC00047\|NCRNA00047\|NEAT2\|PRO2853 | TIA1 | PCG | ENSG00000116001 | TIA-1\|WDM | View Details | expression association |
| RID02610 | MALAT1 | lncRNA | ENSG00000251562 | HCN\|LINC00047\|NCRNA00047\|NEAT2\|PRO2853 | LATS1 | PCG | ENSG00000131023 | WARTS\|wts | View Details | expression association |

D

RG4BP - QUADRatlas

| GENE NAME | BIOTYPE | ENSEMBL ID (RGBP) | GENE ALIAS | CHROMOSOME | START | END | STRAND | KNOWN RBP | RBP TYPE |
|---|---|---|---|---|---|---|---|---|---|
| ELAVL1 | protein_coding | ENSG00000066044.15 | ELAV1\|HUR\|Hua\|MelG | 19 | 7958573 | 8005659 | - | True | RG4-BP |

E

### LncRNA-RNA Interactions - NPInter

| NPINTER INTERACTION ID | INTERACTOR NAME | INTERACTOR TYPE | INTERACTOR ID | TARGET NAME | TARGET TYPE | TARGET ID | INTERACTION MECHANISM | INTERACTION LEVEL | INTERACTION CLASS | INTERACTION DESCRIPTION |
|---|---|---|---|---|---|---|---|---|---|---|
| ncRI-40002126 | MALAT1 | lncRNA | NONHSAG008675 | hsa-mir-101-1 | miRNA | MI0000103 | miRNA target interaction;RNA-RNA interaction | RNA-RNA | regulatory | collections from lncRNome |
| ncRI-40002132 | MALAT1 | lncRNA | NONHSAG008675 | hsa-mir-124 | miRNA | MI0000445 | miRNA target interaction;RNA-RNA interaction | RNA-RNA | regulatory | collections from lncRNome |
| ncRI-40002133 | MALAT1 | lncRNA | NONHSAG008675 | hsa-miR-145-3p | miRNA | MI0000461 | miRNA target interaction;RNA-RNA interaction | RNA-RNA | regulatory | collections from lncRNome |

F

### LncRNA-RNA Interactions LncTarD

| REGULATION ID | REGULATOR NAME | REGULATOR TYPE | REGULATOR ENSEMBLE ID | REGULATOR ALIASES | TARGET NAME | TARGET TYPE | TARGET ENSEMBLE ID | TARGET ALIASES |
|---|---|---|---|---|---|---|---|---|
| RID00070 | MALAT1 | lncRNA | ENSG00000251562 | HCN\|LINC00047\|NCRNA00047\|NEAT2\|PRO2853 | miR-142-3p | miRNA | NA | NA |
| RID00254 | MALAT1 | lncRNA | ENSG00000251562 | HCN\|LINC00047\|NCRNA00047\|NEAT2\|PRO2853 | miR-145 | miRNA | NA | NA |
| RID00375 | MALAT1 | lncRNA | ENSG00000251562 | HCN\|LINC00047\|NCRNA00047\|NEAT2\|PRO2853 | miR-101 | miRNA | NA | NA |

**Figure S4. LncRNA-G4 interacting partner section of CanLncG4.** A) LncRNA search bar. B-C) LncRNA-Protein interactions from B) NPInter, and C) LncTarD. D) Details of RNA G4-binding proteins (RGBP). E-F) LncRNA-RNA interactions from E) NPInter, and F) LncTarD. The red box highlights the tabs that can be used to proceed to the next section.
